# ADAMTS-1 and Syndecan-4 intersect in the regulation of cell migration and angiogenesis

**DOI:** 10.1101/686733

**Authors:** Jordi Lambert, Kate Makin, Sophia Akbareian, Robert Johnson, Stephen D Robinson, Dylan R. Edwards

## Abstract

The extracellular proteoglycanase ADAMTS-1 has critical roles in organogenesis and angiogenesis. We demonstrate here the functional convergence of ADAMTS-1 and the transmembrane heparan sulfate proteoglycan syndecan-4 in influencing adhesion, migration, and angiogenesis *in vitro*. Knockdown of ADAMTS-1 in fibroblasts and endothelial cells resulted in a parallel reduction in cell surface syndecan-4 that was not due to altered syndecan-4 expression or internalization, but was attributable to increased expression and activity of matrix metalloproteinase-9 (MMP-9), a known syndecan-4 sheddase. Knockdown of either syndecan-4 or ADAMTS-1 led to enhanced endothelial cell responses to exogenous vascular endothelial growth factor (VEGF), and increased microvessel sprouting in *ex vivo* aortic ring assays, correlating with reduced ability of the cells to sequester VEGF. On fibronectin but not type 1 collagen matrices, endothelial cells with knockdown of either ADAMTS-1 or syndecan-4 elicited increased migration and showed similarly altered focal adhesion (FA) morphologies, with a higher proportion of larger FA’s and formation of long fibrillar integrin α5-containing FA’s. However, integrin α5-null endothelial cells also displayed enhanced migration in response to ADAMTS-1/syndecan-4 knockdown, indicating that integrin α5 was not the mediator of the altered migratory behaviour. Plating of naïve endothelial cells on cell-conditioned matrix from ADAMTS-1/syndecan-4 knockdown cells demonstrated that the altered behaviour was matrix dependent. Fibulin-1, a known ECM co-factor of ADAMTS-1, was expressed at reduced levels in ADAMTS-1/syndecan-4 knockdown cells. These findings support the notion that ADAMTS-1 and syndecan-4 are functionally interconnected in regulating cell migration and angiogenesis, via the involvement of MMP-9 and fibulin-1 as collaborators

## Introduction

The ADAMTS (a disintegrin and metalloproteinase with thrombospondin motifs) family of extracellular proteases includes 19 members in humans, with diverse roles in tissue development and homeostasis(Kuno *et al*., 1997; Porter *et al*., 2005). Their essential functions are underlined by the recognition that several family members encode genes which are responsible for inherited genetic disorders when mutated, while others are associated with pathologies including cancer, arthritis and cardiovascular disease when aberrantly expressed(Hubmacher and Apte, 2015; Kelwick, Desanlis, *et al*., 2015; Mead and Apte, 2018).

The ADAMTSs are zinc-dependent metalloproteinases with a compound domain structure, each possessing a catalytic domain containing the metalloproteinase and disintegrin-like features, followed by a modular ancillary domain that differs between sub-groups of family members and is important for their biological actions(Kuno *et al*., 1997; Porter *et al*., 2005). The largest clade in the ADAMTS family are identified as proteoglycanases that can cleave a variety of proteoglycans, including versican, aggrecan, brevican, as well as other extracellular matrix proteins(Hubmacher and Apte, 2015; Kelwick, Desanlis, *et al*., 2015). First discovered in 1997, ADAMTS-1 is the prototype of the family and a member of the proteoglycanase clade which also includes ADAMTS-4, -5, -8, -9, -15 and -20. The ability of ADAMTS-1 to cleave structural ECM components is physiologically relevant as demonstrated by *Adamts1-* knock-out mice, which exhibit abnormally high rates of perinatal lethality due to multiple organ defects, in particular severe kidney malformation and cardiac defects (De *et al*., no date; Krampert *et al*., 2005)The surviving female mice suffer from infertility, due to the ineffective cleavage of versican during ovarian maturation(Shindo *et al*., 2000; Mittaz *et al*., 2004; Krampert *et al*., 2005).

However, as well as its proteolytic function, ADAMTS-1 also interacts with other proteins including latent TGF-β(Bourd-Boittin *et al*., 2011) and fibulin-1, which acts as a co-factor(Lee *et al*., 2005). ADAMTS-1 has many context-dependent effects in biological processes such as migration, invasion and cell signalling, which are relevant to its impact on physiology and pathophysiology(De *et al*., no date), indicating it acts through multiple mechanisms. This is reflected in its anti-angiogenic actions, which involve both proteolytic and non-proteolytic mechanisms, the former by mediating the release of highly anti-angiogenic fragments of thrombospondin-1 (TSP1) and TSP-2(Lee *et al*., 2006; Gustavsson *et al*., 2010) and the latter via direct binding and sequestration of vascular endothelial growth factor VEGF_165_ (Luque, Carpizo and Iruela-Arispe, 2003; Fu *et al*., 2011).

Another significant proteoglycan partner of ADAMTS-1 is syndecan-4(Rodríguez-Manzaneque *et al*., 2009). Syndecan-4 is a ubiquitously expressed heparan-sulfate proteoglycan that acts as a key mediator of several cellular processes including adhesion, proliferation and endocytosis(Couchman and Woods, 1999; Elfenbein *et al*., 2012; Elfenbein and Simons, 2013). Its heparan-sulfate glycosaminoglycan (GAG) chains provide binding sites for heparin-binding growth factors such as fibroblast growth factors (FGFs), platelet-derived growth factors (PDGFs) and vascular endothelial growth factors (VEGFs)(Elfenbein and Simons, 2013). The binding of these growth factors to syndecan-4 can have several consequences: activation of cellular signalling can occur through syndecan-4 acting as a co-receptor that presents the growth factor ligand to its signalling receptor, as in the case of FGF, or there can be direct activation of downstream signalling mediated by syndecan-4 itself, such as PKC-α(Oh, Woods and Couchman, 1997b, 1997a). In addition, syndecan-4 can regulate growth factor bioavailability by acting as a cell-bound reservoir that can be released by subsequent proteolytic cleavage(Bergers *et al*., 2000; Ramnath *et al*., 2014). But besides its role as a signalling regulator, syndecan-4 is also a key mediator in focal adhesion formation. Fibroblasts from syndecan-4 null mice exhibit impaired adhesion to fibronectin(Ishiguro *et al*., 2000). Via the binding and activation of protein kinase C α (PKCα), syndecan-4 facilitates α5β1 integrin binding to its substrate fibronectin, allowing maturation of focal adhesions(Mostafavi-Pour *et al*., 2003; Bass *et al*., 2007). Given its key role as a nexus of signalling and adhesion mechanisms, the relative levels and localisation of syndecan-4 are therefore critical determinants of cellular behaviour.

Several reports have connected the actions of ADAMTS enzymes with syndecan-4 (SDC4), including ADAMTS-1 and -4(Rodríguez-Manzaneque *et al*., 2009), ADAMTS-5(Echtermeyer *et al*., 2009; Wang *et al*., 2011), ADAMTS-6 and -10(Cain *et al*., 2016) and ADAMTS-15(Kelwick, Wagstaff, *et al*., 2015). In this study, we have uncovered details of a complex inter-relationship between ADAMTS-1 and syndecan-4 in murine fibroblasts and endothelial cells. We have shown that acute depletion of ADAMTS-1 leads to a concomitant reduction in cell surface levels of syndecan-4, such that down-regulation of either syndecan-4 or ADAMTS-1 have similar consequences on cell behaviour, shown by increases in cellular migration and striking changes to focal adhesions, both of which were fibronectin-dependent. Furthermore, loss of either ADAMTS-1 or syndecan-4 in endothelial cells led to increases in angiogenesis which we demonstrate is due to reduced ability of cells to sequester the pro-angiogenic growth factor VEGF. Moreover, these effects have downstream consequences on α5 integrin trafficking, but α5 integrin is not essential for them to occur. These results demonstrate the existence of an interplay between ADAMTS-1 and syndecan-4 which orchestrates cell adhesion, migration and angiogenesis.

## Results

### Knockdown of ADAMTS-1 results in reduction of cell surface syndecan-4

We have used siRNA to deplete either ADAMTS-1 or Syndecan-4 in murine 3T3 fibroblast cell line (3T3s) and lung microvascular polyoma middle T-antigen immortalised endothelial cells (ECs). mRNA knockdown was confirmed using TaqMan qPCR (Figure S1). Our initial hypothesis was that since ADAMTS-1 has been reported to cleave the N-terminal domain of syndecan-4, depletion of ADAMTS-1 would result in accumulation of syndecan-4 on the cell surface (Rodríguez-Manzaneque *et al*., 2009). However, flow cytometric analysis revealed that siRNA knockdown of ADAMTS-1 resulted in a significant concomitant decrease in cell surface syndecan-4, equivalent to that seen with syndecan-4 knockdown (Figure 1A). To confirm this visually, immunocytochemistry was performed, however as in our hands staining using commercially available syndecan-4 antibody was unsuccessful, cells were transfected with a HA-tagged syndecan-4 construct (HA-SDC4). This confirmed reduced cell surface HA-SDC4 display in ADAMTS-1-depleted cells when compared to cells treated with non-targeting control siRNA (Figure 1B).

**Figure 1:**
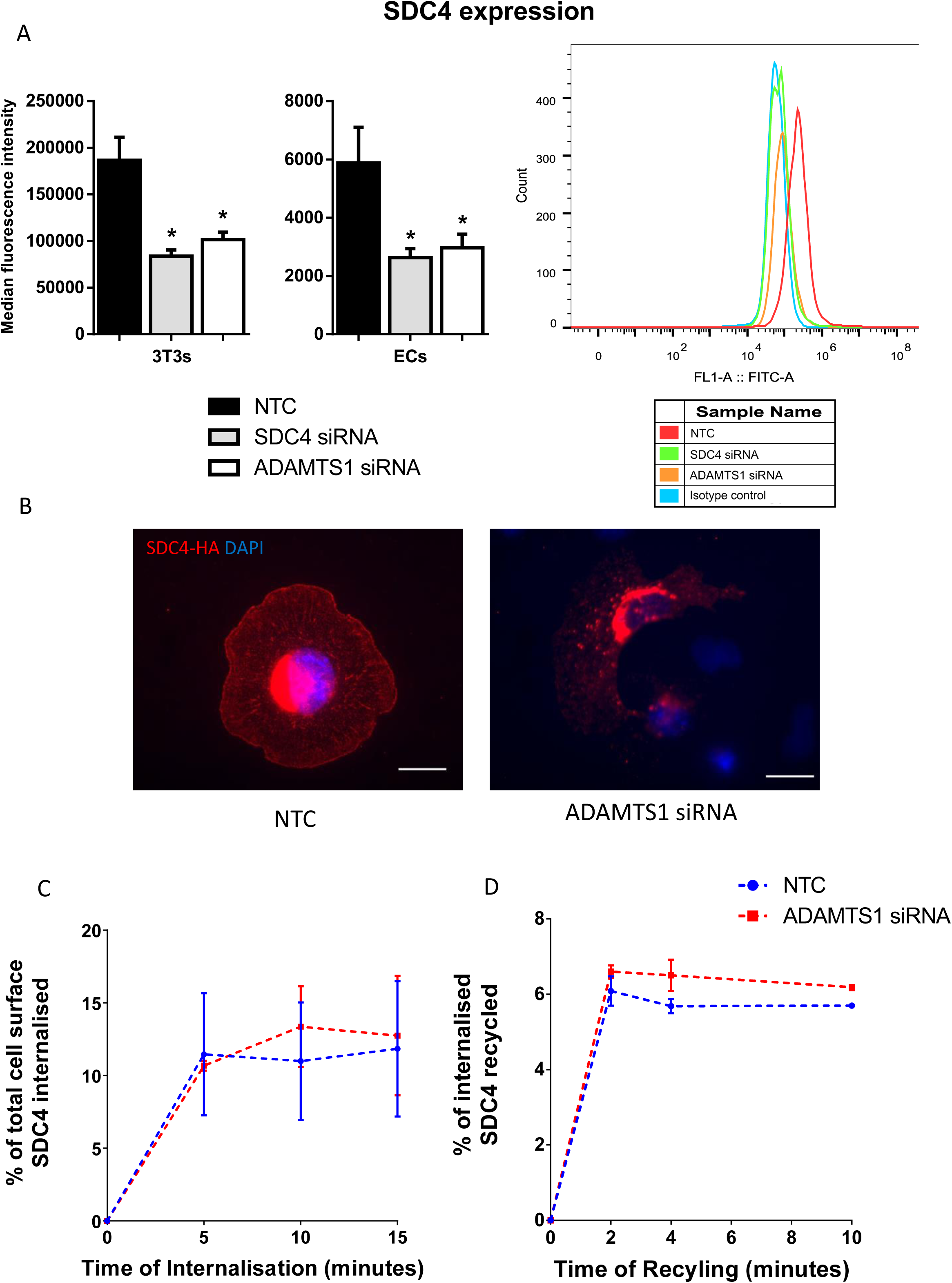
ADAMTS-1 regulates syndecan-4 expression but not via altered recycling or internalisation. **A)** Flow cytometric analysis of syndecan 4 (SDC4) surface levels on lung microvascular endothelial cells (ECs) and 3T3 fibroblast cells (3T3s) treated with non targeting control (NTC), SDC4, or ADAMTS-1 siRNA. The graph (far right) is a representative histogram showing median fluorescence intensity. Values were calculated after gating on forward and side scatter and normalised to isotype controls. N = 3 independent experiments. Error bars represent S.E.M, *P<0.05. **B)** Representative image of HA-tag (*red*) and DAPI (*blue*) immuno-stained ECs expressing HA-tagged SDC4 treated with siRNA against ADAMTS-1 or NTC on fibronectin. N = 3 independent experiments. Scale bar = 20µM. **C)** SDC4 internalisation assay; EC surface proteins were biotinylated using a cleavable, membrane non-permeable biotin. ECs were incubated to allow internalisation, remaining surface biotin was removed using the reducing agent MESNA. A SDC4 capture ELISA was performed on lysed ECs. Streptavidin was used to detect biotinylated SDC4 (N=3, bars represent S.E.M). **D)** SDC4 recycling assay; after the first MesNa treatment, internalised biotin was allowed to return to cell surface. Biotinylated protein that had returned to the surface was again removed using MesNa and an ELISA was performed as in C (N=3, bars represent S.E.M).

To elucidate the mechanism by which ADAMTS-1 regulates syndecan-4 expression, we first investigated whether knockdown of ADAMTS-1 led to changes in *Sdc4* gene expression. ADAMTS-1 siRNA did not alter *Sdc4* expression at the RNA level (Figure s1). Therefore, we hypothesised that loss of cell surface syndecan-4 could be a result of changes in membrane trafficking, so we carried out cell surface biotinylation-based internalization and recycling assays. ECs were surface-labelled with cleavable biotin, incubated for time points to allow internalisation, then biotin remaining on the cell surface was cleaved. Syndecan-4 internalisation was quantified by a syndecan-4 capture ELISA and biotin detection using streptavidin (Figure 1C). For the recycling assay cells were incubated further, to allow biotinylated protein to recycle to the surface (Figure 1D). These experiments demonstrated that neither the rate of syndecan-4 internalization or levels of recycling to the membrane were changed as a result of ADAMTS-1 knockdown.

### Loss of cell surface syndecan-4 is MMP9 dependent

Since reduced syndecan-4 levels did not appear to be a result of altered transcription or membrane trafficking, we speculated that the decreased cell surface syndecan-4 was a result of increased shedding. As MMPs are known cleavers of syndecans they were considered as possible mediators of this shedding(Manon-Jensen, Itoh and Couchman, 2010). ECs were treated with one of the following three broad-spectrum MMP inhibitors: BB-94, CT-1746, GM 6001 or a DMSO vehicle control. Treatment with any of the three MMP inhibitors resulted in accumulation of syndecan-4 at the cell surface, and in cells treated with ADAMTS-1 siRNA, syndecan-4 levels recovered to match that of the NTC cells (Figure 2A). This suggests that MMPs are responsible for the reduction in cell surface syndecan-4 seen in response to loss of ADAMTS-1.

**Figure 2:**
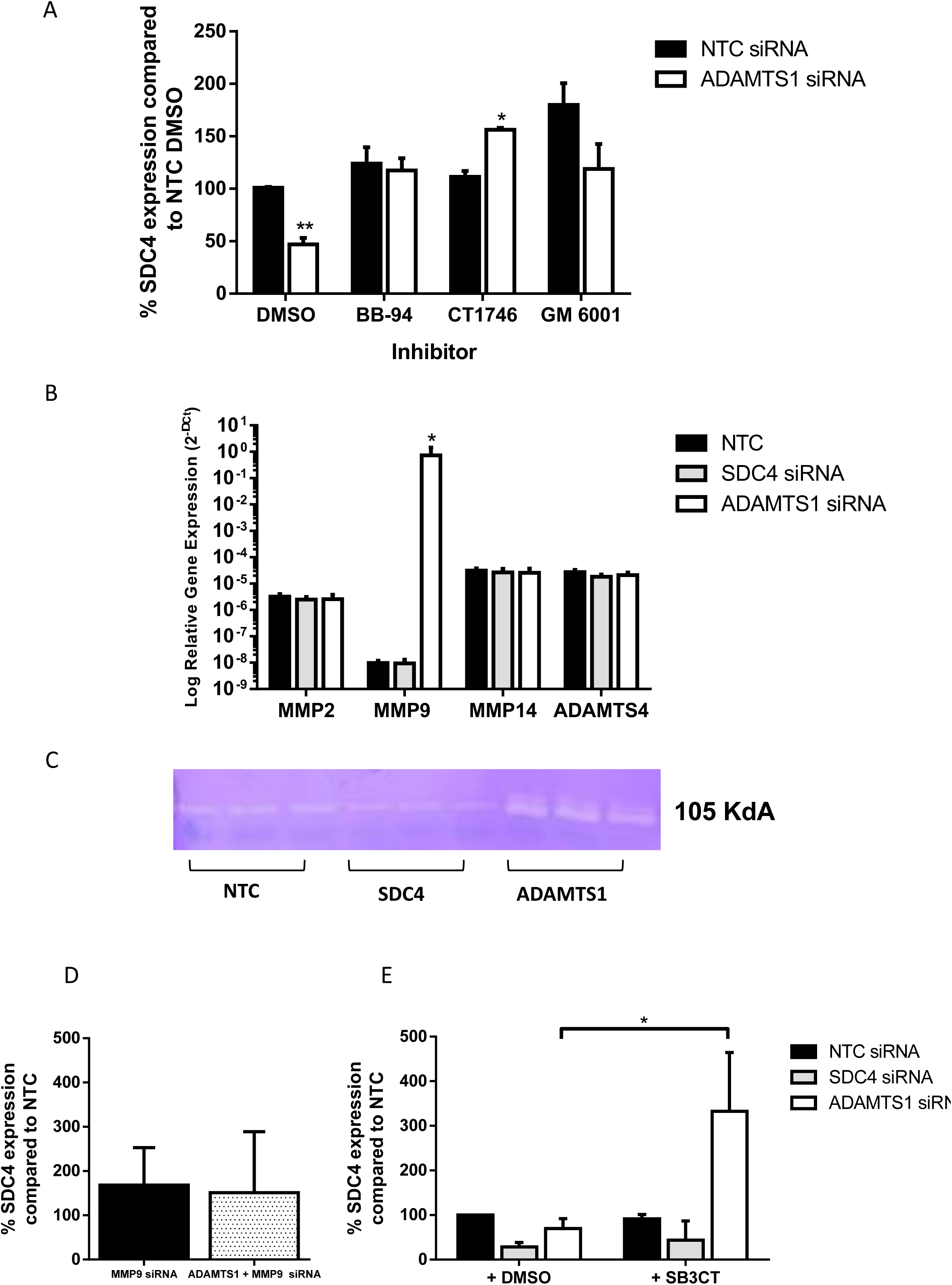
ADAMTS-1 siRNA mediated loss of surface SDC4 is dependent on MMP activity. **A)** Flow cytometric analysis of SDC4 cell surface levels. ECs were transfected with NTC or ADAMTS-1 siRNA, then treated with an MMP inhibitor: BB-94, CT1746, GM 6001 or DMSO vehicle control. Bar chart shows percentage SDC4 expression relative to NTC DMSO treated cells (N=3, *P<0.05, **P<0.01). **B)** TaqMan qPCR for MMP2, MMP9, MMP14 and ADAMTS4 on RNA from ECs treated with NTC, SDC4 or ADAMTS1 siRNA. Bar chart shows relative expression, normalised to a housekeeping control (18s) (N=4, *P<0.05). **C)** A representative gelatin zymogram showing MMP9 activity in the conditioned media of siRNA treated ECs. **D)** Flow cytometric analysis of SDC4 surface levels. ECs were transfected with MMP9, or a combination of MMP9 and ADAMTS-1 siRNA. Bar chart shows percentage SDC4 expression relative to NTC siRNA treated cells. **E)**

To identify the specific enzyme responsible, a panel of MMPs known to be expressed in endothelial cells were profiled: MMP2, MMP9, MMP14 (MT1-MMP) and ADAMTS-4(Bergers *et al*., 2000; Genís *et al*., 2006; Hsu *et al*., 2012). ADAMTS-1 siRNA knockdown resulted in significantly increased expression of MMP9 compared to NTC or syndecan-4 siRNA treated cells, which expressed almost no MMP9. In both treated and untreated cells, MMP2, MMP14 and ADAMTS-4 expression was unchanged (Figure 2B). The increase in MMP9 RNA expression was reflected in increased MMP9 activity, as shown by gelatin zymography of conditioned media from ADAMTS-1 siRNA, syndecan-4 siRNA and NTC treated cells (Figure 2C). In these experiments we were unable to detect syndecan-4 that had been shed from the cell surface into the media under any conditions, indicating that this is likely rapidly degraded. However treatment with MMP9 siRNA (Figure 2D) or an MMP9 specific inhibitor SB-3CT (Figure 2E) was sufficient to reverse the phenotype resulting from ADAMTS-1 knockdown. This demonstrates that increased expression of MMP9, known to cleave syndecan-4 is directly responsible for the reduced syndecan-4 levels seen after ADAMTS-1 siRNA treatment(Ramnath *et al*., 2014; Reine *et al*., 2019).

### Syndecan-4 contributes to ADAMTS-1’s sequestration of VEGF

ADAMTS-1 acts as an inhibitor of angiogenesis by sequestering VEGF165, a key pro-angiogenic factor. This sequestration prevents VEGF from binding and activating its major receptor VEGFR2(Luque, Carpizo and Iruela-Arispe, 2003). ADAMTS-1 exclusively binds VEGF165, and the interaction is dependent on the heparin binding domain. Previous work by Iruela-Irispe *et al* has suggested that a heparin sulphate proteoglycan such as a member of the syndecan family may provide a bridge to facilitate this interaction(IRUELA-ARISPE, CARPIZO and LUQUE, 2003).

The impact of ADAMTS-1 or syndecan-4 siRNA on VEGF signalling was investigated. Endothelial cells were serum-starved for 3 hours, then stimulated with 30ng/mL VEGF164 (the mouse equivalent of human VEGF165). Cells were lysed at 0, 5, 15 and 30 minutes post stimulation and lysates collected were run on a Western blot to look for changes in the VEGF signalling pathway. Upon siRNA knockdown of either ADAMTS-1 or syndecan-4, increased phospho-VEGFR2 and phospho-ERK were seen, indicating an increase in cellular responsiveness to exogenous VEGF when ADAMTS-1 or syndecan-4 levels are reduced (Figure 3A, figure S2).

**Figure 3:**
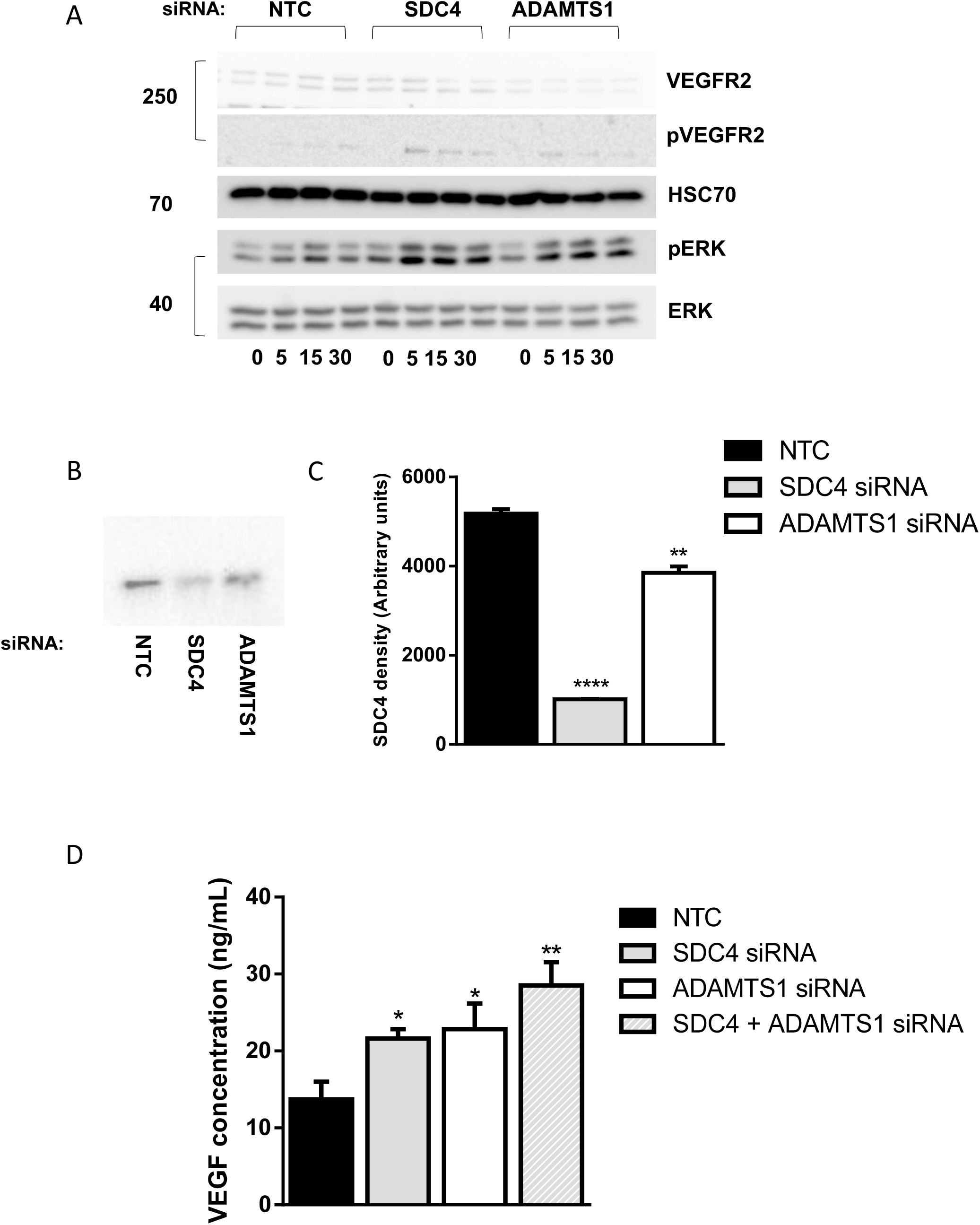
SDC4 contributes to the sequestration of VEGF164 by ADAMTS-1. **A)** VEGF signalling time-course. siRNA-treated ECs were adhered overnight on fibronectin. Serum-starved cells were stimulated with 30 ng/mL VEGF for 0, 5, 15 and 30 minutes. Western blots were performed on lysates with anti-VEGFR2, -ERK, and their phosphorylated forms. HSC70 was used as a loading control. Blot is representative of 4 independent experiments. **B)** Western blot using an anti-SDC4 antibody on fractions from VEGF immunoprecipitations (IPs) carried out on HUVECs with lentiviral silencing targeting NTC, ADAMTS-1 or SDC4. **C)** Image J densitometric quantification of IP western blots (N=3, error bars represent S.E.M, **P<0.01, ****P<0.0001). **D)** VEGF sequestration ELISA on ECs treated with NTC, SDC4, or ADAMTS-1 siRNA. Cells were kept at 4°C, incubated in serum free media containing 30 ng/mL VEGF 164 for 30 minutes. Media was collected and a VEGF sandwich ELISA was performed to detect unbound VEGF (N=3, bars represent S.E.M, *P<0.05, **P<0.01).

In order to test the hypothesis that syndecan-4 may be aiding ADAMTS-1’s sequestration of VEGF, co-immunoprecipitations were performed. An IP for VEGF was able to co-precipitate syndecan-4, revealing that syndecan-4 either directly or indirectly binds VEGF. When cells were treated with ADAMTS-1 siRNA, less syndecan-4 was found upon co-precipitation (Figure 3B, 3C). The reduction in syndecan-4 seen with ADAMTS-1 siRNA may be an indicator that the interaction involves ADAMTS-1, however it may also be a result of the reduced cell surface syndecan-4 seen with ADAMTS-1 siRNA treatment.

As both ADAMTS-1 and syndecan-4 are capable of binding VEGF, the relevance of these interactions was investigated using a VEGF ELISA to assess how much VEGF cells were able to sequester. As before, 30 ng/mL VEGF was added to ECs cooled to 4°C to prevent signalling. A Western blot was performed to confirm that signalling pathways were not activated (figure S2A), and qPCR confirmed VEGF RNA levels were unchanged (figure S2B). The cells were kept at 4°C for 30 minutes to allow binding. The media was then collected and an ELISA was performed to quantify the amount of unbound VEGF remaining in the media. When cells were treated with ADAMTS-1, syndecan-4 or both siRNAs, the amount of free VEGF remaining in the media significantly increased (Figure 3D). Taken together then, these data suggest that the loss of ADAMTS-1 or syndecan-4 prevents the sequestration of VEGF, leaving it freely available to activate downstream signalling pathways.

### ADAMTS-1 and syndecan-4 sequestration of VEGF inhibits angiogenesis

The physiological relevance of the ability of ADAMTS-1 and syndecan-4 to sequester VEGF was next investigated. The process of angiogenesis requires both proliferation and migration of endothelial cells, which can be stimulated by VEGF. BrdU labelling of siRNA treated cells revealed increased proliferation in ADAMTS-1-depleted cells, but not syndecan-4-depleted (Figure 4A). As syndecan-4 is known to interact with numerous other growth factors (FGFs, EGFs, PDGFs) this may be a result of multiple pathway involvement(Harburger and Calderwood, 2008).

**Figure 4:**
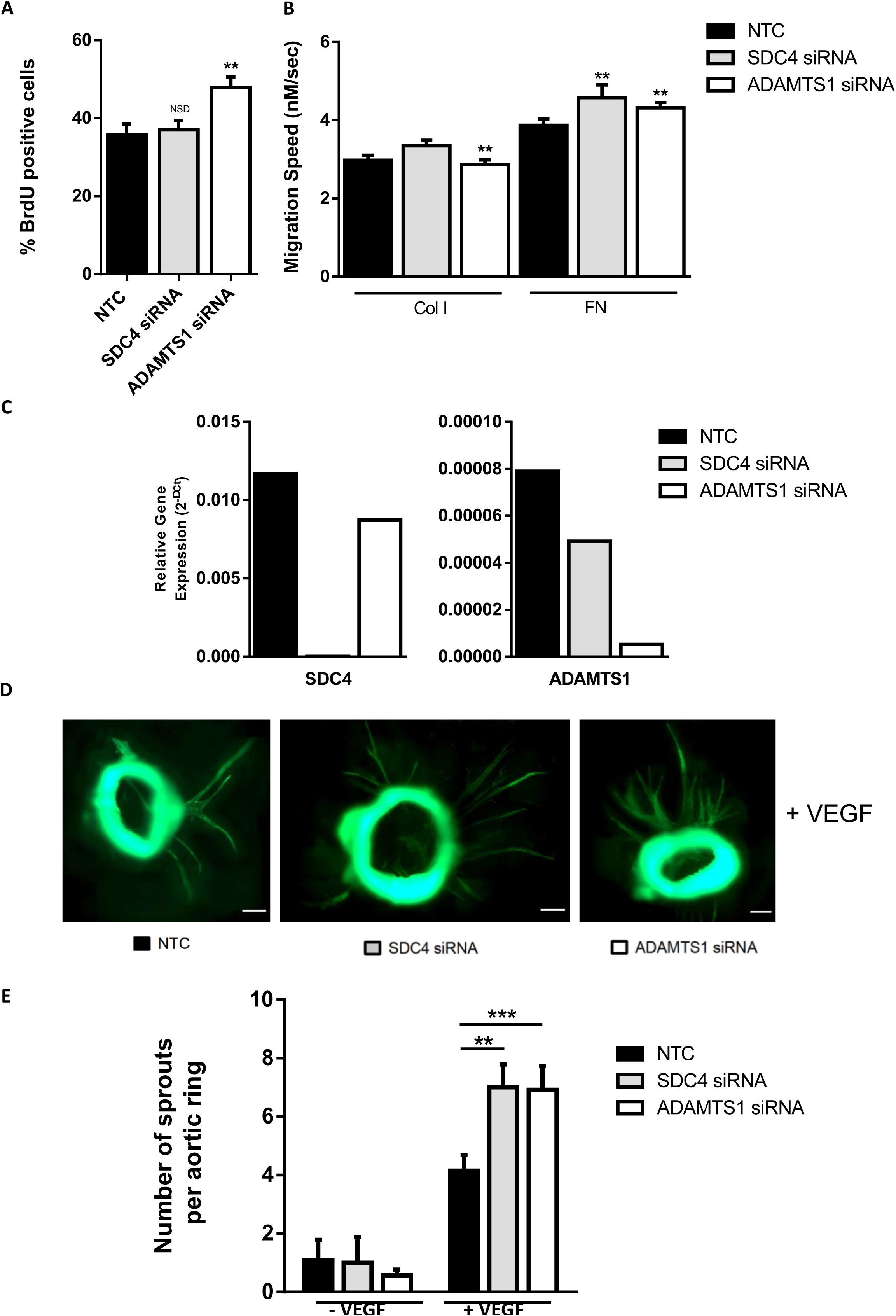
ADAMTS-1 and SDC4 siRNA treatment results in increased angiogenesis. **A)** Proliferation assay on siRNA-treated ECs allowed to incorporate BrdU for 12 hours. Cells were fixed and stained for BrdU and DAPI. Total number of nuclei and BrdU positive nuclei were counted, bar chart shows the percentage of proliferating cells (N=4, **P<0.01, ns = not significant). **B)** 2D migration assay on siRNA treated ECs seeded onto 10 µg/mL collagen I or fibronectin matrix and allowed to adhere overnight. Migration velocity was determined with time lapse microscopy over a period of 16 hours (N= ≥100 cells in 4 independent experiments, error bars represent S.E.M, **P<0.001). **C)** Taqman qPCR of aortic rings treated with ADAMTS1 or SDC4 siRNA showing relative levels of SDC4 and ADAMTS-1 in 3 independent experiments, bars represent S.E.M, **p<0.001**. D)** Micro vessel sprouting of aortic ring explants from 6-8 week old mice treated with indicated siRNA. Pictures are representative images of aortic rings stained with FITC-conjugated BS1-lectin, scale bar = 200 µm. **E)** Bar chart showing the total number of micro vessel sprouts per aortic ring 6 days post-VEGF-stimulation (n≥50 rings per condition, N = 4 independent experiments, error bars represent S.E.M, **P<0.001, ***P<0.0001).

As conflicting reports exist with regards to the roles of ADAMTS-1 and syndecan-4 in cell migration(Krampert *et al*., 2005; Bass *et al*., 2007; Rodríguez-Manzaneque *et al*., 2009; Obika *et al*., 2012), random migration assays were carried out on siRNA-depleted cells to assess effects of their knockdown on motility. siRNA-treated endothelial cells were plated on either 10 µg/mL collagen I or 10 µg/mL fibronectin and time-lapse microscopy was performed over a period of 16 hours to track cell migration. Migration speed was significantly increased with both ADAMTS-1 and syndecan-4 on fibronectin matrices only (Figure 4B).

To see if the altered proliferation and migration resulted in a corresponding increase in angiogenesis, the mouse aortic ring assay was utilised. Aortas were harvested from 6-8 week-old mice, sliced into rings and treated with siRNA against ADAMTS-1 or syndecan-4. The rings were embedded in matrix, and the number of sprouts that formed was used as a marker of angiogenesis(Baker *et al*., 2011). Both siRNAs successfully depleted their targets in the aortic rings (Figure 4C). Both ADAMTS-1 and syndecan-4 depletion resulted in a marked increase in VEGF-dependent new vessel sprouting (Figure 4D, 4E), demonstrating a physiologically relevant effect of the sequestration of VEGF by ADAMTS-1 and syndecan-4.

### Adhesions are altered in ADAMTS-1/syndecan-4 siRNA cells on fibronectin matrices

As knockdown of either ADAMTS-1 or syndecan-4 led to altered migration speeds on fibronectin but not on collagen I, focal adhesions parameters were assessed on both matrices. Cells were seeded on a collagen I or fibronectin matrix for 90 or 180 minutes. Staining for paxillin was performed to visually assess focal adhesions, and their size and number was calculated using ImageJ. While on both collagen and fibronectin the numbers of focal adhesions were unchanged in siRNA-depleted cells, adhesions to fibronectin matured more quickly. At 90 minutes post-adherence to fibronectin, the average focal adhesion size was larger, with a higher percentage of adhesions reaching a more mature stage (Figure 5A, Figure 5B). The majority of adhesions remained as small nascent adhesions (<2 µm^2^) (NTC: 95.6%, syndecan-4: 90.9%, ADAMTS-1: 92%), however a higher percentage of cell adhesions in siRNA-treated cells had developed into focal adhesions (2-6 µm^2^) (NTC: 4.3%, syndecan-4: 8.5%, ADAMTS-1: 7.2%). By 180 minutes this difference no longer existed, suggesting that the siRNA-treated cells do not form more large adhesions, rather their adhesions mature at a faster rate. Adhesions to collagen were unaffected (Figure S4), confirming that the observed effects are due to interaction with the fibronectin matrix. These changes in focal adhesions of ADAMTS-1 or syndecan-4 knockdown cells on fibronectin were also apparent in a marked increase in paxillin signalling in response to VEGF (Figure 5C, figure S5). This supports the proposition that ADAMTS-1 and syndecan-4 act to sequester VEGF reducing its bioavailability, and that their depletion leads to enhanced VEGF signalling and subsequent changes to focal adhesion morphology.

**Figure 5:**
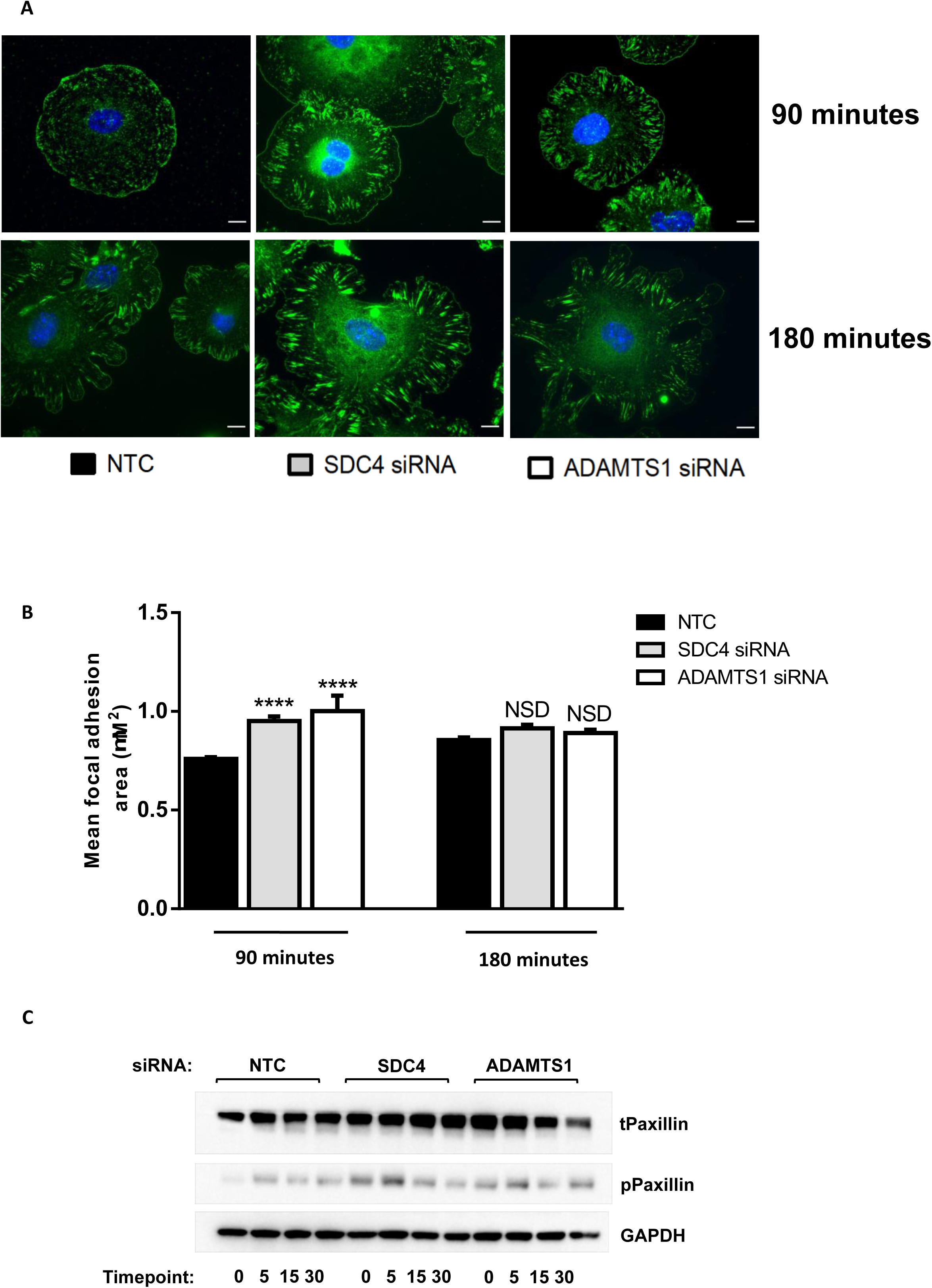
siRNA depletion of ADAMTS-1 or SDC4 results in altered focal adhesions. **A)** Representative images of siRNA-treated ECs adhered to fibronectin coated glass coverslips for 90 or 180 minutes. Cells fixed and stained for focal adhesions with paxillin (*green*) and the nucleus with DAPI (*blue*). N =3 independent experiments. Scale bar = 10 µm. **B)** Area of paxillin positive focal adhesions, calculated using ImageJ. Graph shows average size of adhesions, N=20 cells per conditions from 3 independent experiments, error bars represent SEM, ****P<0.00001. **C)** Western blot analysis of paxillin signalling using an anti phospho-paxillin antibody in siRNA-treated ECs adhered to fibronectin, serum starved for 3 hours then stimulated with 30ng/mL VEGF for 0, 5 15 and 30 minutes. GAPDH was used as a loading control. N = 3 independent eaxperiments.

### Alpha 5 integrin localisation is disrupted in the absence of ADAMTS-1 or syndecan-4

Since our data showed perturbation of fibronectin-dependent adhesions following ADAMTS-1 or syndecan-4 depletion, the contribution of alpha 5 integrin (α5), the major fibronectin receptor in nascent focal adhesions in endothelial cells, was considered. Immunocytochemical visualisation of α5 revealed the development of long, fibrillar-like adhesions after 180 minutes adherence to fibronectin in cells treated with ADAMTS-1 or syndecan-4 siRNA, as apposed to the more traditional α5 localisation in focal adhesions <6 µm^2^ seen in NTC cells (Figure 6A).

**Figure 6:**
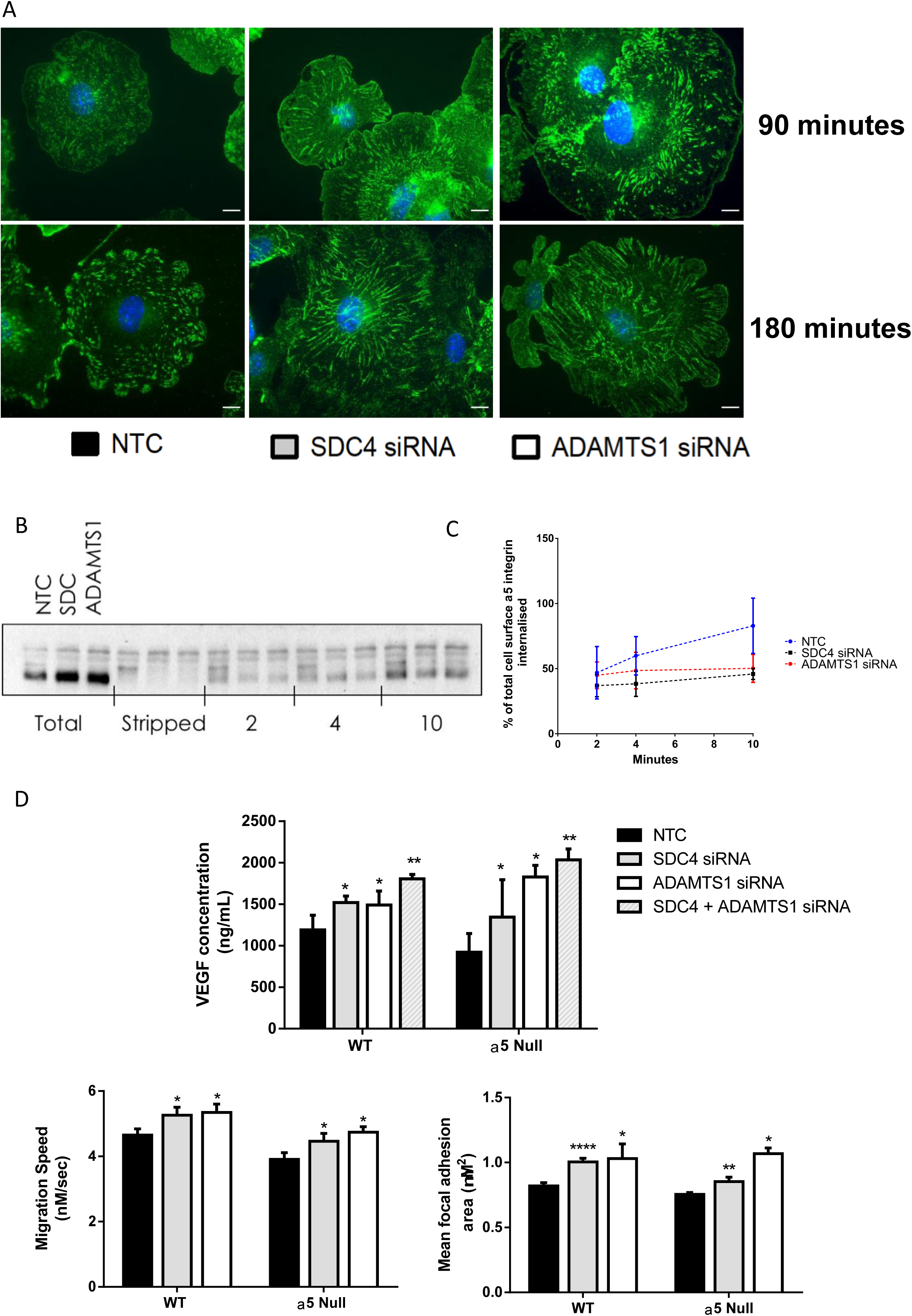
Depletion of ADAMTS-1 or SDC4 results in downstream alterations to alpha5 integrin behaviour. **A)** Representative images of siRNA-treated ECs adhered to fibronectin coated glass coverslips for 90 or 180 minutes. Cells fixed and stained for α5 integrin (*green*) and DAPI (*blue*). N = 3 independent experiments. Scale bar =10 µm. **B)**Western blot of IP lysates from internalisation assays using an anti-alpha5 integrin antibody. Cell-surface proteins of ECs seeded on fibronectin were biotinylated, post-internalisation at 2, 4 and 10 minutes. Remaining surface biotin was ‘stripped’ using a membrane-impermeable reducing agent (MeSna). Cells were lysed and immunoprecipitated (IP’d) for biotin. The blot shows ‘total’ surface alpha5 (from cells not internalised and not stripped), cells not internalised, but ‘stripped’ as controls, and alpha5 internalisation at each time point. N = 3 independent experiments. **C)** Quantification of alpha5 integrin internalisation assays **D)** Experiments from 3D, 4B, and 5C were repeated in ECs isolated from α5 null mice and their wildtype controls. (N=3 independent experiments, error bars represent S.E.M, *P<0.05, **P<0.01, ****P<0.00001).

As syndecan-4 is known to have a role in integrin surface trafficking, biotinylation-based internalisation assays were performed as before to assess α5 membrane trafficking(Morgan *et al*., 2013). To quantify internalisation, immunoprecipitation of biotin followed by Western blot detection of α5 integrin was utilised. Western blotting revealed an increase in total cell surface α5 in ADAMTS-1 and syndecan-4 siRNA treated cells, as well as decreased levels of internalisation, suggesting the altered adhesions may be a result of α5 accumulation due to impaired internalisation (Figure 6B, figure 6C). To test whether this retention of α5 integrin on the cell surface in ADAMTS-1/ syndecan-4 knockdown cells was responsible for the altered cell characteristics we again performed the experiments above in α5-null endothelial cells treated with ADAMTS-1 or syndecan-4 siRNA. However, although α5-null endothelial cells showed reduced migration compared to their wild-type counterparts, the same phenotypes of increased rates of migration and focal adhesion maturation, and decreased VEGF sequestration following ADAMTS-1 or syndecan-4 knockdown were maintained (Figure 6D). These data suggest that the changes in α5 integrin seen are likely a downstream consequence of changes in cell behaviour mediated by ADAMTS-1/ syndecan-4 knockdown, rather than α5 integrin being the key effector of the phenotypes.

### EC behaviour and signalling phenotypes are maintained in response to conditioned matrix

As the mechanism by which ADAMTS-1 and syndecan-4 siRNA increased cell migration and changed focal adhesions was not dependent on integrin α5, modulation of the ECM itself was considered, especially since these responses were reliant on the cells being on a fibronectin matrix. In order to assess this, experiments were performed using ‘conditioned matrix’ (CM) from siRNA-treated cells that were seeded onto glass coverslips and allowed to produce matrix for 48 hours, following which the siRNA-treated cells were then stripped away, leaving behind CM. Untreated cells were then seeded onto the CM and left to adhere for 3 hours and immunocytochemistry for α5 integrin was carried out. The phenotype of long, fibrillar adhesions was repeated, indicating that α5 integrin changes occur in response to a modified matrix (Figure 7A). Signalling was examined in a similar fashion: untreated cells were adhered to CM for 45, 90 and 180 minutes, then cells were lysed and signalling responses examined. Cells adhered to the conditioned matrix from ADAMTS-1 or syndecan-4 knockdown cells showed increased paxillin and ERK signalling, suggesting that the altered matrix is sufficient to induce the more migratory phenotype (Figure 7B).

**Figure 7:**
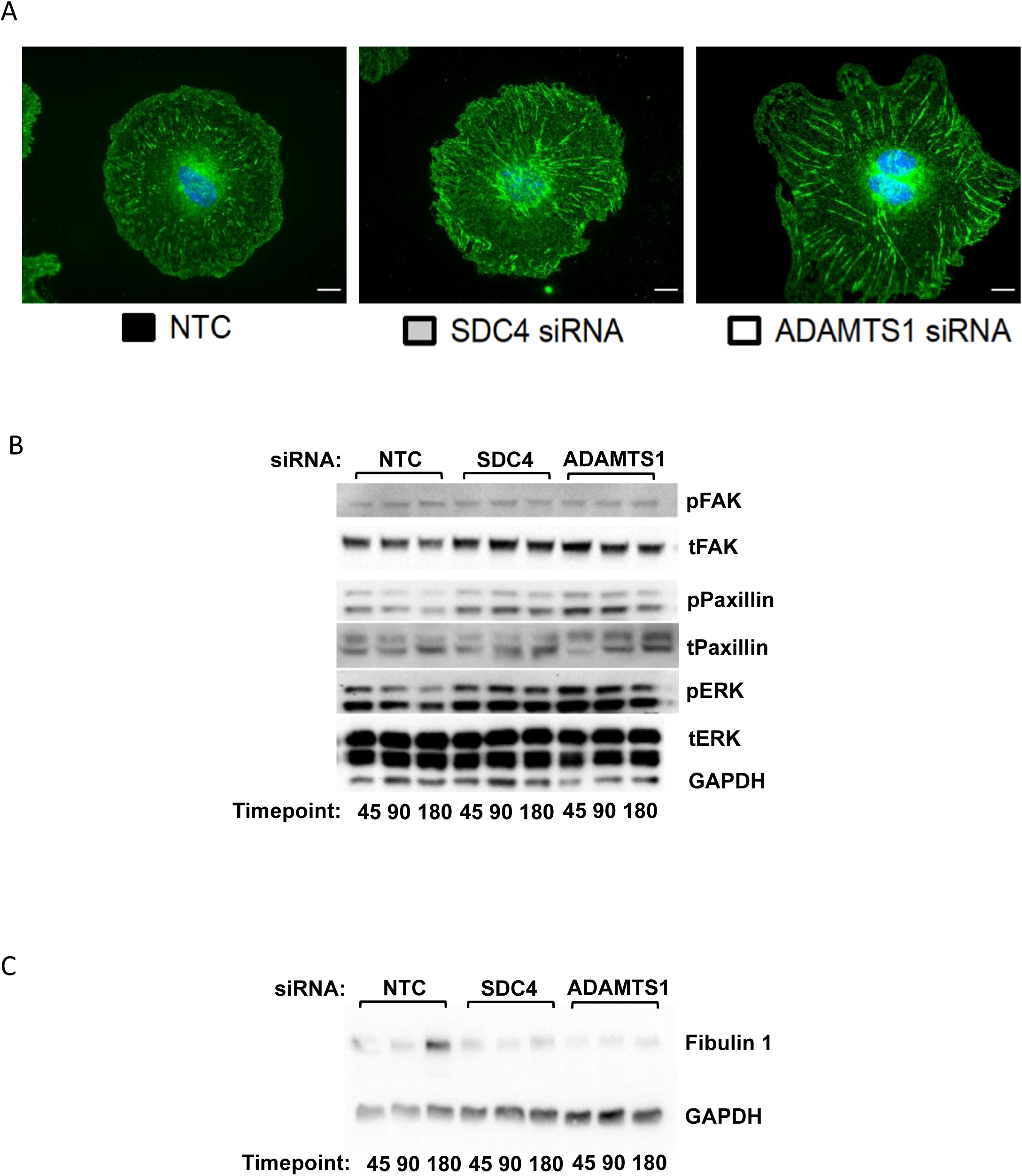
EC behaviour and signalling phenotypes are maintained in response to conditioned matrix. **A)** Representative images of untreated ECs after 180 minutes on conditioned matrix (CM) from siRNA-treated ECs allowed to produce matrix for 48 hours before being removed. Cells were fixed and stained for alpha5 integrin (*green*). N =3 independent experiments. Scale bar = 10 µm. **B)** Western blot of untreated ECs seeded on CM for 45, 90 and 180 minutes, using anti-total (t) and phosphorylated(p) FAK, ERK and Paxillin. GAPDH was used as a loading control. N=3 independent experiments**. C)**Western blot for fibulin 1 of siRNA treated ECs seeded on fibronectin for 45, 90 or 180 minutes. GAPDH was used as a loading control. N=3 independent experiments

Fibulin 1, a co-factor for ADAMTS-1 and a matrix protein which has been reported to inhibit migration showed enhanced expressed at 180 minutes post adhesion to fibronectin in NTC treated cells, but not ADAMTS-1 or syndecan-4 siRNA treated cells (Figure 7C, figure S7)(Twal *et al*., 2001; Lee *et al*., 2005). These data support a mechanism by which ADAMTS-1 and syndecan-4 regulate adhesion by inducing expression of the matrix protein fibulin 1.

## Discussion

Our work provides new insights into a functional connection between ADAMTS-1 and the cell surface proteoglycan syndecan-4. Knockdown of either protein via siRNA results in similar responses with regards to angiogenesis, cell migration, and integrin α5 trafficking. Our data support the notion that ADAMTS-1 is an inhibitor of angiogenesis, and we offer novel data that this occurs in co-ordination with syndecan-4.

Previous work has shown that ADAMTS-1 inhibits angiogenesis through direct sequestration of the major pro-angiogenic growth factor VEGF165, reducing its bioavailability(Luque, Carpizo and Iruela-Arispe, 2003). This has also been shown to be true for ADAMTS-4, which can bind VEGF and inhibit angiogenesis in a similar fashion to ADAMTS-1(Hsu *et al*., 2012). The heparin binding domain of VEGF165 is necessary for its interaction with ADAMTS-1, and work by Iruela-Arispe *et al* has suggested that this interaction may be reliant on a heparin-like molecule. One such candidate is syndecan-4, a heparan sulphate proteoglycan and the major syndecan in endothelial cells, known to interact with heparin-binding growth factors such as VEGFs, PDGFs and FGFs(Harburger and Calderwood, 2008). Thus, given the numerous connections between ADAMTS-1 and syndecan-4 and the dependence of ADAMTS-15 on syndecan-4 to enhance migration ability, we sought to investigate the relationship between syndecan-4 and ADAMTS-1.

Importantly, ADAMTS-1 has been reported to cleave the ectodomain of syndecan-4, promoting migration in a way that reflects genetic deletion of the proteoglycan(Rodríguez-Manzaneque *et al*., 2009). However, we were unable to detect evidence of a cleaved form of syndecan-4 released from the fibroblasts and endothelial cells used in our studies. Instead we were able to demonstrate a convergent functional relationship between ADAMTS-1 and syndecan-4 since ADAMTS-1 siRNA treatment caused a significant reduction in syndecan-4 display at the cell surface, indicating that syndecan-4 is dependent on ADAMTS-1. A key finding from our work is that this connection between ADAMTS-1 and cell surface expression of syndecan-4 involves matrix metalloproteinase-9 (MMP-9).

Syndecan ectodomain shedding is an important regulatory mechanism allowing for rapid changes in cell surface receptor dynamics; shedding occurs juxtamembrane and can be carried out by a number of matrix proteinases(Manon-Jensen, Itoh and Couchman, 2010). MMP7 and 14 are both able to shed the extracellular domain of syndecan-2 and MMP-9 has been shown to shed syndecan-4 from the cell surface in response to TNFα(Kwon *et al*., 2014; Ramnath *et al*., 2014; Lee *et al*., 2017). There was a significant increase in MMP9 expression in ADAMTS-1 siRNA-treated cells, and upon loss of MMP9, via either an inhibitor or siRNA-depletion, syndecan-4 was no longer lost from the cell surface. We therefore propose that the loss of ADAMTS-1 results in increased shedding of syndecan-4 that is mediated by MMP9.

At this point, we are not able to identify the mechanism by which ADAMTS-1 knockdown results in increased MMP9 expression. One possibility however is that it occurs as a result of a shift in the angiogenic signalling balance in endothelial cells. ADAMTS-1 is anti-angiogenic, whereas MMP9 is highly pro-angiogenic, and can trigger the ‘angiogenic switch’, a process in tumours where the balance of pro and anti-angiogenic factors swings towards a pro-angiogenic outcome(Bergers *et al*., 2000). MMP9 promotes release of VEGF bound in the matrix to heparan-sulphate chains through cleavage(Hawinkels *et al*., 2008). VEGF induces MMP9 expression which leads to elevated free VEGF levels resulting in a positive feedback loop(Hollborn *et al*., 2007). Similarly, VEGF can upregulate ADAMTS-1 expression - suggesting a negative feedback mechanism at play(Xu, Yu and Duh, 2006). It seems possible that loss of ADAMTS-1 disturbs the fine balance of VEGF signalling, resulting in the cells switching to a more angiogenic phenotype, and the upregulation of MMP9.

We sought to investigate the possibility that syndecan-4 works co-operatively with ADAMTS-1 to inhibit angiogenesis through sequestration of VEGF. This idea contrasts with several hypotheses that syndecan-4 acts as a co-receptor for VEGF to enhance signalling, as is the case for FGFs. Through Co-IPs, we demonstrated that syndecan-4 does indeed bind VEGF. We interpret this binding as sequestration since an increase in VEGF signalling and free VEGF is seen upon syndecan-4 depletion. These data strongly suggest that syndecan-4 does not promote VEGF signalling, but instead alongside ADAMTS-1 sequesters VEGF to inhibit angiogenesis. We further demonstrated this finding in a physiologically relevant *ex vivo* model of angiogenesis and observed increased micro-vessel sprouting in an aortic ring assay with ADAMTS-1 or syndecan-4 siRNA-depletion. It therefore seems plausible that MMP9, ADAMTS-1 and syndecan-4 work together in a complex equilibrium to balance VEGF bioavailability and regulate angiogenesis.

For angiogenesis to occur, endothelial cells must proliferate and migrate - processes that are heavily influenced by VEGF signalling. Loss of ADAMTS-1 resulted in increased proliferation, however this result was not mirrored with loss of syndecan-4. This implies that the two molecules have distinct as well as shared actions. On the one hand, syndecan-4 is a major signalling nexus involved as a partner in the actions of several growth factors as well as a source of intracellular signals itself. These multiple pathways could negate the impact of syndecan-4’s sequestration of VEGF. Likewise, ADAMTS-1 functions independently of syndecan-4, specifically in relation to angiogenesis by releasing antiangiogenic peptides from TSP-1 and –2 TSP-2 through cleavage(Lee *et al*., 2006). At the organism level, although *Sdc4* and *Adamts1* null mice share a delayed wound healing response(Echtermeyer *et al*., 2001), the models do not phenocopy each other. Whilst *Sdc4* knockout mice are relatively healthy, *Adamts1* KO mice exhibit developmental issues with stunted growth and high embryonic mortality(Shindo *et al*., 2000; Echtermeyer *et al*., 2001).

Previous work on fibroblasts from *Sdc4*-/- animals showed syndecan-4 was necessary for directional migration to occur(Bass *et al*., 2007), however in our experiments, acute depletion of syndecan-4 or ADAMTS-1 in endothelial cells resulted in increased migration speed. This observation was only seen on fibronectin, with decreased or no change in migration speed on collagen. Fibronectin matrix-dependent migration effects have been noted previously with the ADAMTSs; work by Kelwick *et al* showed that the closely related ADAMTS-15 was able to inhibit MDA-MB-231 cell migration on fibronectin matrices, and that this effect was dependent on syndecan-4(Kelwick, Wagstaff, *et al*., 2015). In endothelial cells, α5 integrin functions as the major fibronectin receptor. Interestingly, syndecan-4 regulates adhesion synergistically with fibronectin and α5β1 integrin(Morgan, Humphries and Bass, 2007), as well as regulating integrin trafficking. Phosphorylation of syndecan-4 acts as a molecular switch, controlling levels of αVβ3 and α5β1 internalisation and recycling, and affecting cell behaviour(Morgan *et al*., 2013).

Due to the clear connection between syndecan-4, fibronectin, and integrins, we sought to investigate α5 integrin as a potential mediator of migratory effects. Immunocytochemical and cell surface biotinylation data support an important role for syndecan-4 in regulation of α5 integrin trafficking. ADAMTS-1 knockdown also appeared to regulate α5 integrin trafficking, but whether this is a direct effect is unclear, though it would be consistent with the resulting loss of syndecan-4. Despite this, siRNA knockdowns of ADAMTS-1 and syndecan-4 repeated in α5 null endothelial cells maintained the phenotypes of increased VEGF signalling, increased free VEGF, faster random migration and larger focal adhesions, suggesting that α5 integrin is not necessary for ADAMTS-1 and syndecan-4 regulated angiogenesis and migration. Therefore, we hypothesised that knockdowns of ADAMTS-1 and syndecan-4 may result in ECM alterations, resulting in downstream changes to integrin trafficking. Both proteins interact with the ECM: ADAMTS-1 is secreted and anchors in the ECM while syndecan-4 forms cell-ECM attachments(Kuno and Matsushima, 1998; Gopal *et al*., 2017). In support of this, non-treated naïve cells plated on ‘conditioned matrix’ matrix created by ADAMTS-1/ syndecan-4 siRNA cells behaved in a similar manner to siRNA treated cells, forming altered α5 adhesions, and increased paxillin signalling.

Fibulin 1 is an ECM protein, and a co-factor for ADAMTS-1(Lee *et al*., 2005); expression of fibulin 1 has been reported to result in a fibronectin-specific inhibition of migration in a cell type-dependent manner(Twal *et al*., 2001). In fibroblasts, this effect is dependent on syndecan-4(Williams and Schwarzbauer, 2009). Fibulin 1 is indeed upregulated at 180 minutes post adhesion in NTC-treated cells, but not in ADAMTS-1 or syndecan-4 siRNA cells. We therefore propose a mechanism for the control of migration in endothelial cells where ADAMTS-1/ syndecan-4 regulate production of fibulin 1, thereby suppressing migration and facilitating α5 integrin turnover.

This work builds on the current literature on ADAMTS-1 as an anti-angiogenic protein; demonstrating co-operation between ADAMTS-1 and the heparin sulphate proteoglycan syndecan-4. We highlight a role for syndecan-4 in inhibiting migration and angiogenesis, which contrasts with some previous published data. When attempting to understand conflicts with regards to syndecan-4 it is important to consider the model used. Much previous work has focused on the constitutive knockout mouse, and cells and tissues isolated from it(Echtermeyer *et al*., 2001). Compensatory mechanisms from developmental adaptation to loss of syndecan-4 are possible(Bass *et al*., 2007). However it is now recognised that passenger mutations in knockout mice created using embryonic stem cells from the 129 strain, may confound interpretation of data from these mice(Eisener-Dorman, Lawrence and Bolivar, 2009). There is a potential argument that previous studies had used fibroblasts derived from *Sdc4*-knockout mice, which may be differently affected by loss of syndecan-4 than the VEGF-responsive endothelial cells that are the focus of our work, due to the involvement of syndecan-4 in multiple growth factor signalling pathways. However, we have seen similar effects of siRNA-mediated knockdown of syndecan-4 on migration in fibroblasts as in endothelial cells, suggesting these are not dependent on cell type.

In conclusion, our work shows that ADAMTS-1 regulates cell-surface expression of syndecan-4, and in endothelial cells both molecules act via sequestration of VEGF and inhibition of angiogenesis. ADAMTS-1 influences syndecan-4 levels on the cell surface via regulation of MMP9 production. Our studies also reveal convergent roles for ADAMTS-1 and syndecan-4 in regulating the ECM, inhibiting migration, modulating integrin adhesions and activating the matrix protein fibulin 1.

## Materials and methods

### Cell culture and isolation

Human umbilical vein endothelial cells (HUVECs) were cultured in EBM-2 media supplemented with the SingleQuotsTM kit (Lonza, Slough, UK). 3T3 fibroblasts were cultured in high glucose DMEM (invitrogen) supplemented with 10% fetal bovine serum (FBS) (HyClone, Invitrogen), 100 units/mL penicillin/streptomycin (Invitrogen), and 2 mM glutamax (Invitrogen).

Primary mouse lung endothelial cells were isolated from adult mice as previously described by Reynolds & Hodivala-Dilke(Reynolds and Hodivala-Dilke, 2006). For immortalisation, ECs were treated with polyoma-middle-T-antigen (PyMT) retroviral transfection as described by Robinson *et al*(Robinson *et al*., 2009). Immortalized mouse lung endothelial cells (ECs) were cultured in IMMLEC media, a 1:1 mix of Ham’s F-12:DMEM medium (low glucose) supplemented with 10% FBS, 100 units/mL penicillin/streptomycin, 2 mM glutamax, 50 μg/mL heparin (Sigma).

All cells were cultured at 37°C in a humidified chamber with 5% CO_2_. For experimental analyses, plates and flasks were coated with either: human plasma fibronectin (FN) (Millipore), collagen I (Col I) (Fisher Scientific) overnight at 4°C. VEGF-A 164 (henceforth referred to as VEGF) was made in-house according to the method published by Krilleke *et al*(Krilleke *et al*., 2007).

### RNAi and lentiviral transfections

For siRNA transfections, ECs were transfected with 50nM siRNA (Thermo Scientific) using the Amaxa nucleofector system (Lonza) according to manufacturer’s instructions, with a final siRNA concentration of 50nM, and the nucleofection program T-005. For syndecan-4 (SDC4) ICC and ELISA, ECs were transfected with HA-tagged SDC4 construct (a generous gift from James Whiteford). Transfected cells were selected for using a GFP reporter tag.

For shRNA knockdown of syndecan-4/ADAMTS-1 in HUVECs, packaging plasmids (Addgene) were mixed with shRNA plasmid (Mission shRNA, Sigma-Aldrich) in Optimem medium (Invitrogen) and Lipofectamine 2000 (Invitrogen) with the following ratios: 750 ng psPAX2, 250 ng pMD2.G, 1 μg shRNA. The mixture was transferred to 80% confluent 293T cells in 10cm dishes for 6 h. The medium was replaced with regular DMEM 10% FBS and collected after 48 h. Medium containing virus was filtered through a 0.45 μm filter and used for HUVEC transduction with the addition of 8 μg/mL polybrene (Sigma). The target sequences used were as follows:

Human *Sdc4* sequence:

5’-CCGGCCTGATCCTACTGCTCATGTACTCGAGTACATGAGCAGTAGGATCAGGTTT TTG-3’.

Human *Adamts1* sequence:

5’-CCGGCCACAGGAACTGGAAGCATAACTCGAGTTATGCTTCCAGTTCCTGTGGTTT TTG-3’.

### Flow cytometry

For flow-cytometric analysis, cells were removed from culture plates using citric saline buffer (1.35 M KCL, 0.15 M Na3C6H5O7). Cells were collected by centrifugation and resuspended in FACS buffer (5% FBS in PBS) and labelled with anti-mouse syndecan-4 (KY/8.2, BD Bioscience), or isotype control (Mouse IgG2a, Invitrogen) for 1 hour. Cells were washed, resuspended in FACS buffer and incubated with fluorophore-conjugated secondary antibody (eBioscience). Data was collected using a Beckman CytoFLEX and analysed using FlowJo.

For treatment with MMP inhibitors one of: 5 μM BB-94 (Abcam), 10 μM CT1746 (CellTech), 10 μM GM-6001 (Millipore) in 10 μL DMSO, or DMSO vehicle control, was added to cells for 18 hours prior to flow cytometric analysis.

### RNA extraction, reverse transcription–PCR and real-time quantitative RT–PCR analyses

Total cell RNA was extracted using the SV Total RNA Isolation Kit (Promega) according to manufacturer’s instructions. RNA samples were reverse transcribed using GoScript™ Reverse Transcriptase system (Promega). Quantitative real-time TaqMan PCR was carried out as described previously(López-Otín and Matrisian, 2007).

### Immunocytochemistry

siRNA-transfected ECs were seeded at a density of 2×10^5^ cells/well in 24 well plates on acid-washed and oven-sterilised glass coverslips pre-coated with FN or Col I. Cells were fixed at indicated time points in 4% paraformaldehyde, washed in PBS, blocked and permeabilised with 0.3% Triton X-100, 10% serum, and incubated with primary antibody diluted 1:100 in PBS for 1 h at RT. Primary antibodies were: anti-HA tag (Thermo, 2-2.2.14), anti-paxillin (ab32084; Abcam), anti-α5 integrin (Abcam, ab150361). Coverslips were PBS washed, and incubated with the relevant Alexa-Fluor®-conjugated secondary antibody (Invitrogen) diluted 1:500 in PBS for 45 min at RT. Coverslips were washed in PBS again, before mounting on slides with Prolong® Gold containing DAPI (Invitrogen). Focal adhesion area calculations were carried out in FIJI™.

### Internalisation and Recycling assays

#### Internalisation

1×10^6^ siRNA transfected cells were seeded in 10 cm dishes. After adhering overnight, cells were serum starved for 3 hours, transferred to ice, washed twice in cold PBS, and surface labelled at 4°C with 0.3 mg/ml NHS-SS-biotin (Pierce) for 30 min. Labelled cells were washed in cold PBS and transferred to IMMLEC at 37°C to allow internalization. At each time point, medium was removed, dishes were transferred to ice and washed twice with ice-cold PBS. Biotin was removed from proteins remaining at the cell surface by incubation with the membrane impermeable reducing agent MesNa (20 mM MesNa in 50 mM Tris-HCL (pH 8.6)) for 1 hour at 4°C. MesNa was quenched by the addition of 20 mM iodoacetamide (IAA) for 10 mins. Cells were lysed in 200 mM NaCl, 75 mM Tris, 15 mM NaF, 1.5 mM Na_3_VO_4_, 7.5 mM EDTA, 7.5 mM EGTA, 1.5% Triton X-100, supplemented with protease inhibitor (Merck). Lysates were cleared by centrifugation at 10,000 × g for 10 min. Levels of syndecan-4internalisation were determined by capture ELISA. For α5 integrin internalisation analysis, biotinylated protein was isolated by immunoprecipitation using anti-biotin (Jackson Immunoresearch), and levels of α5 integrin assessed by SDS-PAGE.

#### Recycling

After surface labelling, cells were incubated in IMMLEC at 37°C for 20 minutes to allow internalization. Following removal of biotin from surface proteins using MesNA, the internalized fraction was then allowed to recycle to the membrane by returning cells to 37°C in IMMLEC. At the indicated times, cells were returned to ice and biotin was removed from recycled proteins by a second reduction with MesNa. Biotinylated syndecan-4 was then determined by capture-ELISA.

#### Capture-ELISA

96-well microplates (R&D Systems) were coated overnight with 5 μg/mL HA-tag antibody (51064, Proteintech) in PBS at room temperature. The plates were blocked in PBS containing 0.05% Tween-20 (0.05% PBS-T) with 1% BSA for 1 hr at room temperature. HA-SDC4 was captured by 2 hour incubation of 100 μl cell lysate (1 μg/μL) at room temperature. Unbound material was removed by extensive washing with 0.05% PBS-T. Wells were incubated with streptavidin-conjugated horseradish peroxidase (R&D Systems) in 0.05% PBS-T containing 1% BSA for 1 hr at room temperature. Following further washing, biotinylated SCD4 was detected with Tetramethylbenzidine (R&D systems).

### Zymography

Zymography was performed using SDS-PAGE (7.5%) gels co-polymerized with 1 mg/ml gelatin. Protein concentration of media samples were equalized, and added to 5X non-reducing sample buffer (Thermo Scientific). After electrophoresis, gels were washed twice for 30 minutes in wash buffer (2.5% Triton X-100, 50 mM Tris HCl, 5 mM CaCl_2_, 1 μM ZnCl_2_) at room temperature. Gels were incubated overnight in incubation buffer (1% Triton X-100, 50 mM Tris-HCl, 5 mM CaCl_2_, 1 µM ZnCl_2_) at 37°C. Gels were stained in Coomassie Blue R 250 (Thermo Fisher) in a mixture of methanol:acetic acid:water (4:1:5) for 1 hour and destained in the same solution without dye. Gelatinase activities were visualized as distinct bands.

### Western blot analysis

ECs were seeded at 2.5×10^5^ cells per well in 6 well plates coated with 10 μg/ml FN. After 24 hours, cells were starved for 3 hours in serum free medium (OptiMEM®; Invitrogen). VEGF was then added to a final concentration of 30 ng/ml. Cells were lysed at the indicated times (see main text) in RIPA buffer (25mM Tris, pH 7.5, 150 mM NaCl, 0.1% SDS, 0.5% sodium deoxycholate, 1% Triton X-100, supplemented with protease inhibitor (Merck). After protein quantification using the DC BioRad assay, 30 μg of protein from each sample was loaded onto 8% polyacrylamide gels. For paxillin analysis, samples were loaded onto a 4-12% gradient gel for better resolution. The protein was transferred to a nitrocellulose membrane and incubated for 1 h in 5% milk powder/PBS plus 0.1% Tween-20 (0.1% PBS-T), followed by an overnight incubation in primary antibody diluted 1:1000 in 5% bovine serum albumin (BSA)/ 0.1% PBS-T) at 4°C. The blots were then washed 3x with 0.1% PBS-T and incubated with the relevant horseradish peroxidase (HRP)-conjugated secondary antibody (Dako) diluted 1:2000 in 5% milk/PBST, for 1 h at room temperature. Chemiluminescence was detected on a Bio-Rad Gel Doc XR + (Bio-Rad).

Antibodies (all used at 1:1000 and purchased from Cell Signaling Technology, unless noted otherwise) were: anti-phospho (Y1175) VEGFR2 (clone 19A10); anti-VEGFR-2 (clone 55B11); anti-phospho (Thr202/Tyr204) p44/42 MAPK Erk1/2 (clone D13.14.4E); anti-total p44/42 MAPK Erk1/2, anti-HSC70 (clone B-6, Santa Cruz Biotechnology), anti-human syndecan-4 (ab24511, abcam), anti-phospho (Tyr118) paxillin (2541), anti-phospho (Tyr925) FAK, anti-FAK, anti-GAPDH (6C5,abcam).

FIJI™ was used for quantification of band densities.

### Immunoprecipitation assay

HUVECs were grown to 80-90% confluency in 10-cm dishes coated with 10 μg/ml FN in PBS. Cells were lysed in RIPA buffer. 400 μg of total protein from each sample was immunoprecipitated by incubation with protein-G Dynabeads® (Invitrogen) coupled to a mouse-anti-human VEGF-A antibody (VG-1, Abcam) on a rotator overnight at 4°C. Immunoprecipitated complexes were washed three times with 0.2 ml of RIPA buffer, and once in PBS, before being added to, and boiled in NuPAGE® sample reducing agent and sample buffer (Life Technologies) for Western blotting.

### VEGF ELISA

For quantification of free VEGF present in the media, ECs were plated at a density of 2.5×10^5^ cells per well of a 6 well plate. Cells were chilled to 4°C, media was removed and cells were washed twice in ice cold PBS. Media was replaced with ice cold OptiMEM® containing VEGF at a final concentration of 30 ng/ml. Media was diluted 1:10 in 1% BSA in PBS, and VEGF concentration was quantified using a mouse VEGF duo-set ELISA, according to manufacturer’s instructions (DY493, R&D systems).

### Proliferation Assay

siRNA-transfected ECs were seeded on 10 μg/ml FN coated glass coverslips (1.5×10^4^ cells/well of a 24 well plate). After 4 hours, the media was replaced with IMMLEC containing 10nM BrdU. After 12 hours cells were fixed in 4% PFA. To stain, cells were incubated in 1M HCL for 30 mins at room temperature, then permeabilized with PBS 0.25% triton X-100 for 10 minutes and blocked by a 20 minute incubation in DAKO block (Agilent). BrdU was detected by incubation with anti-BrdU (ICR1, Abcam) diluted 1:100 in PBS for 1 hour at room temperature. Coverslips were washed in PBS, then incubated with anti-sheep Alexa-Fluor® (Invitrogen) for 1 hour at room temperature. After further PBS washes, coverslips were mounted in pro-long gold® containing DAPI (invitrogen).

### Random-migration assay

siRNA-transfected ECs were trypsinised and seeded at 1.5×10^4^ cells/well in 24-well plates coated with 10 μg/ml FN or 10 μg/ml Col I in PBS, and allowed to adhere overnight. The media was then replaced fresh IMMLEC. One phase contrast image/well was taken live every 10 min in a fixed field of view using an inverted Axiovert (Zeiss) microscope for 16 h at 37°C and 5% CO_2_. Individual cells were then manually tracked using the FIJI™ cell tracking plugin, mTrackJ and the speed of random migration was calculated in nm moved/seconds.

### Ex vivo aortic ring assay

Thoracic aortae were isolated from 6 to 8 week-old adult mice and prepared for culture as described extensively by Baker *et al*(Baker *et al*., 2011). Syndecan-4 or ADAMTS-1 depletion was induced using 1 μM siRNA and oligofectamine. Where indicated, VEGF was added at 30 ng/mL. Microvessel growth of aortic rings was quantified after 6-10 days.

### Conditioned matrix generation

For experiments using conditioned matrix (CM), siRNA treated cells were seeded at ∼70% confluence and allowed to produce matrix for 48 hours. Plates were washed in PBS and cells were removed by incubation in 20 mM Ammonium Hydroxide. After extensive washing in PBS, untreated cells could then be seeded onto the CM.

## Acknowledgments

We would like to thank Dr. James Whiteford for providing the HA-SDC4 construct. We would also like to thank Andrew Loveday and Robert Hindmarsh, for keeping the lab up and running, Paul Thomas for his microscopy expertise, and all the members of the Robinson Lab. This work was funded by the BBSRC doctoral training partnership.

## Author contributions

J.L. designed and performed experiments. S.D.R. provided the ECs and mice. R.J. bred genotyped and harvested mice. J.L. and K.M. wrote the manuscript. All authors assisted with manuscript review.

---

**Figure S1:**
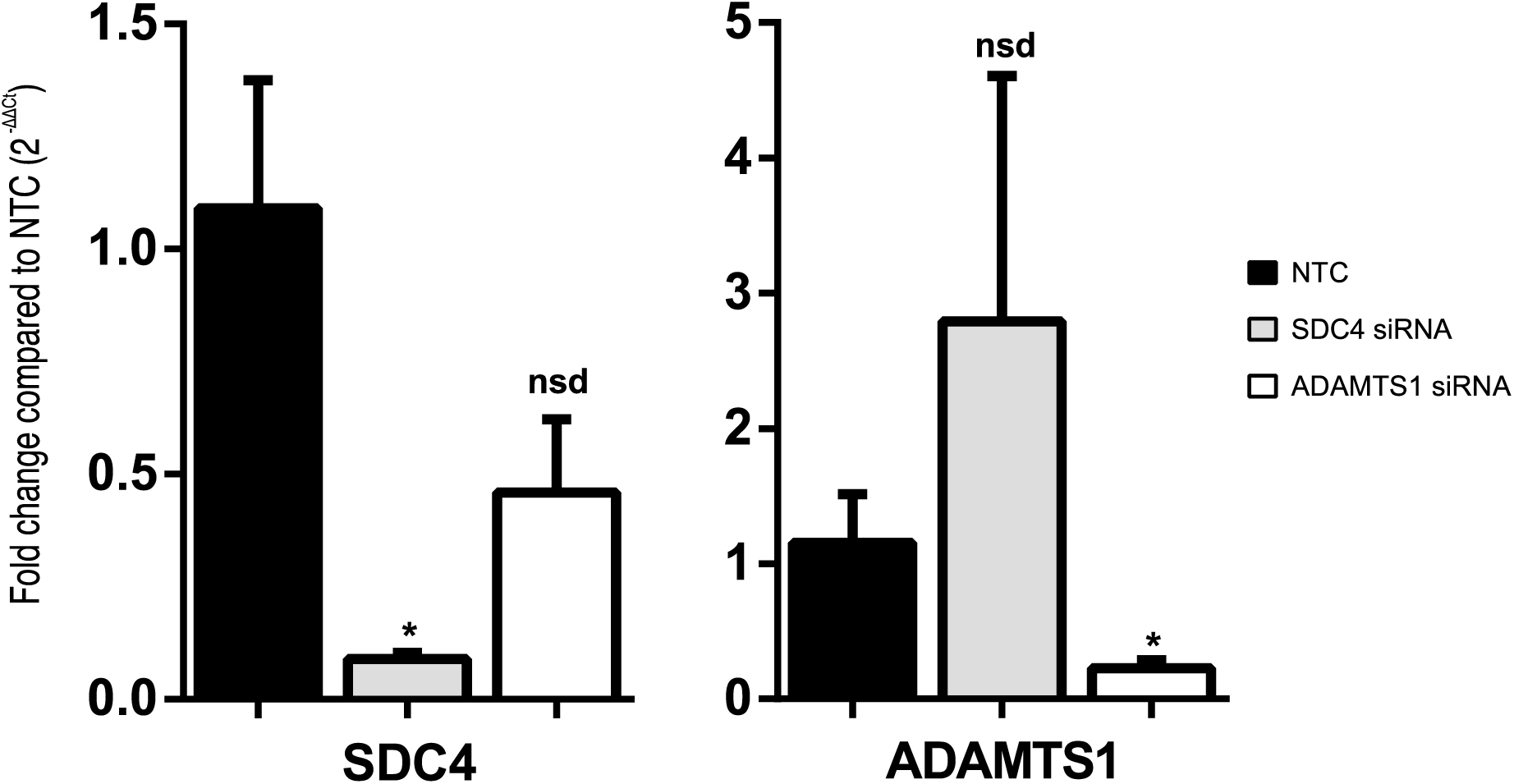
SiRNA successfully depletes targets in endothelial cell lines. Taqman qPCR of Ecs treated with ADAMTS-1 or SDC4 siRNA showing relative levels of SDC4 and ADAMTS-1 in 3 independent experiments, bars represent S.E.M, *p<0.005

**Figure S2:**
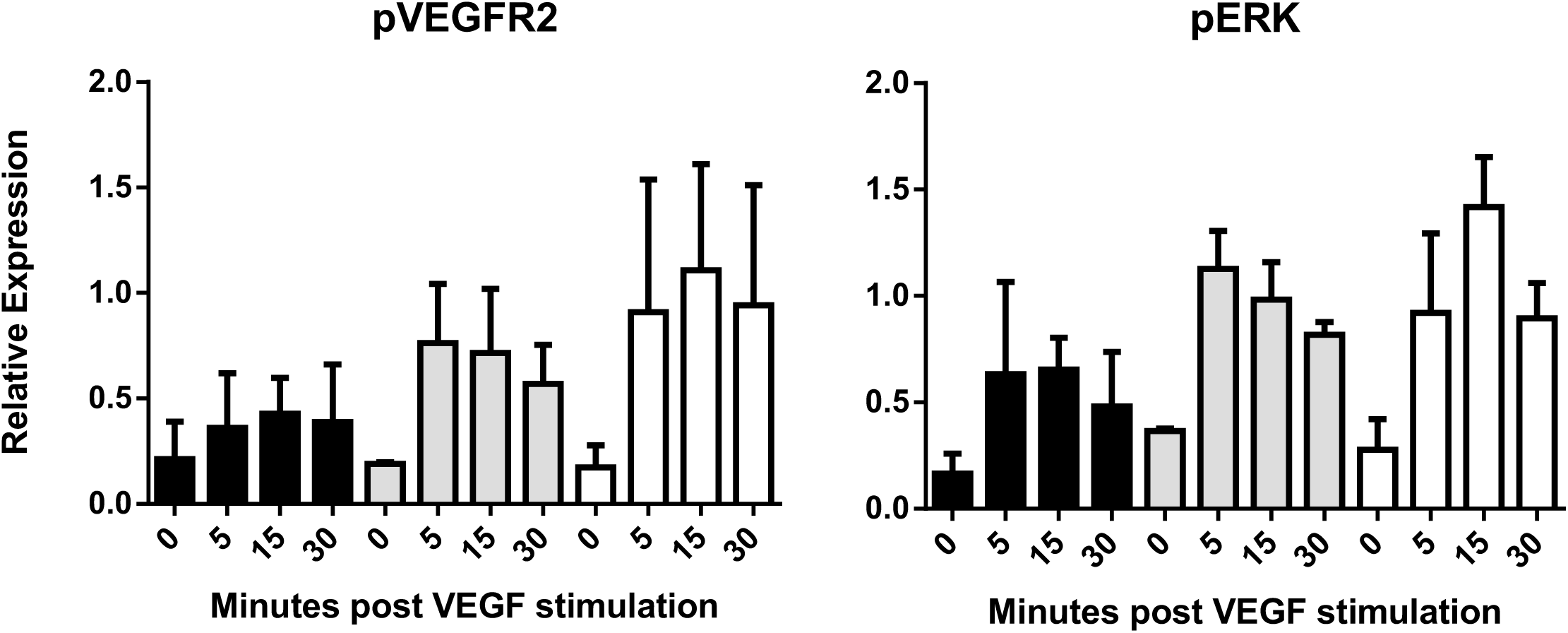
VEGF signalling is increased upon depletion of ADAMTS-1 or SDC4. Image J densitometric quantification of VEGF timecourses (figure 3A). Phosphorylated protein expression levels were calculated relative to total protein. (N=3, error bars represent S.E.M).

**Figure S3:**
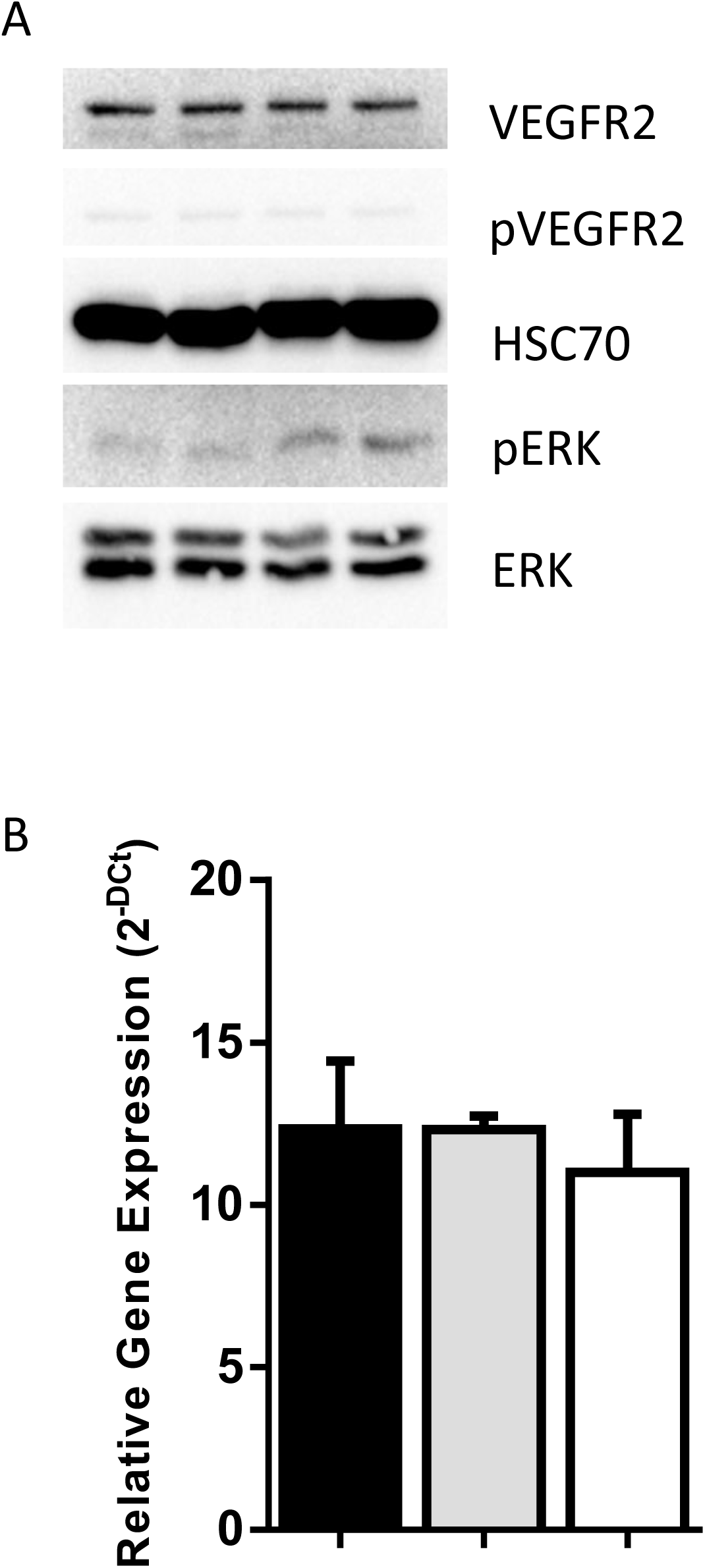
Increased free VEGF in media is not due to increased transcription or altered signalling. **A)** 30 ng/µl VEGF was added to ECs at 4°C, after 30 minutes cells were lysed and western blotted for VEGFR2, ERK and their phosphylated forms. **B)** RNA was collected 48 post siRNA transfection of Ecs, qPCR was performed to determine VEGF expression levels.

**Figure S4:**
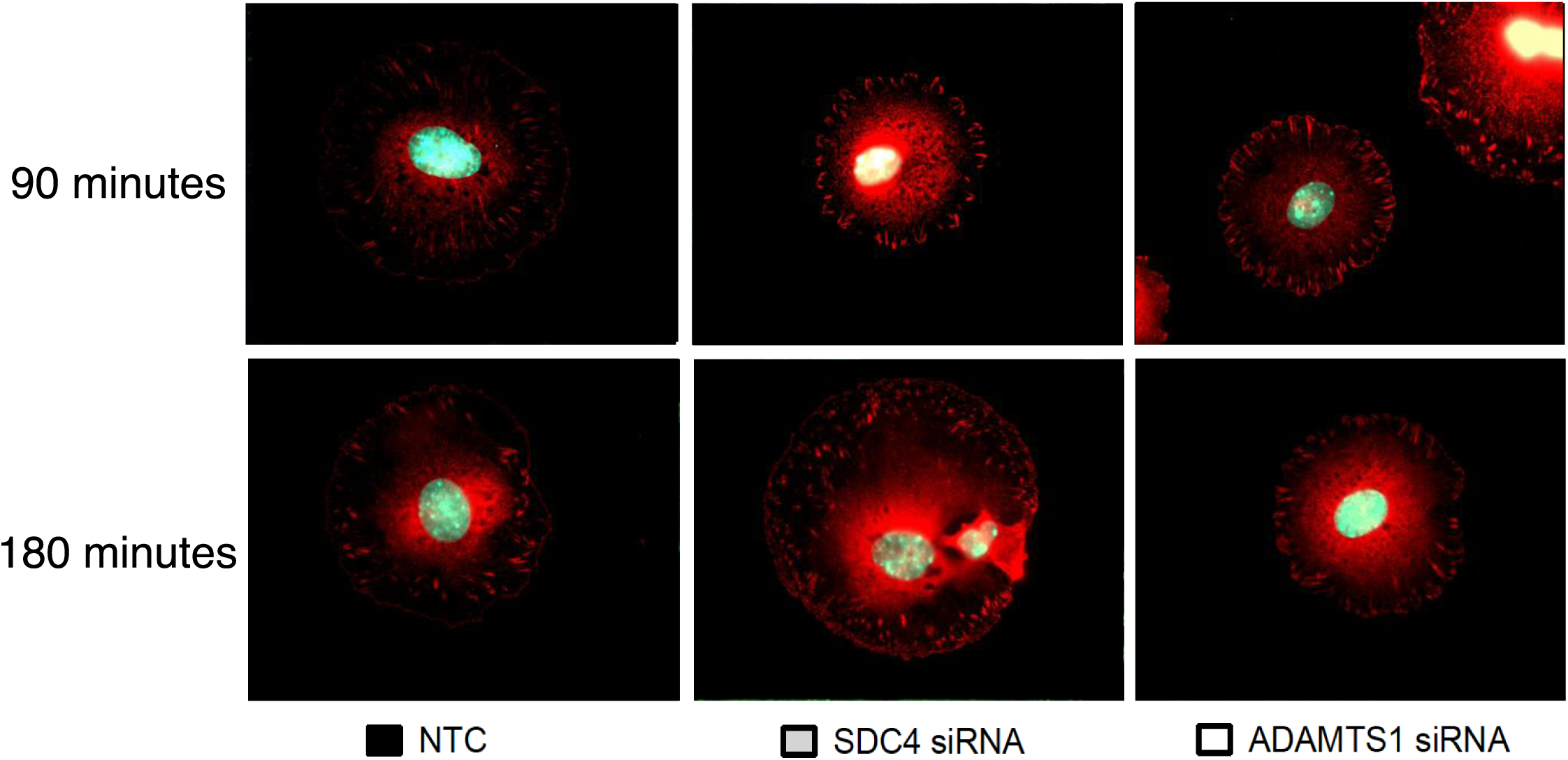
ADAMTS-1 or SDC4 depletion does not affect adhesions to collagen. Representative images of siRNA-treated ECs adhered to collagen coated glass coverslips for 90 or 180 minutes. Cells fixed and stained for focal adhesions with paxillin (*green*) and the nucleus with DAPI (*blue*). N =3 independent experiments. Scale bar = 10 µm.

**Figure S5:**
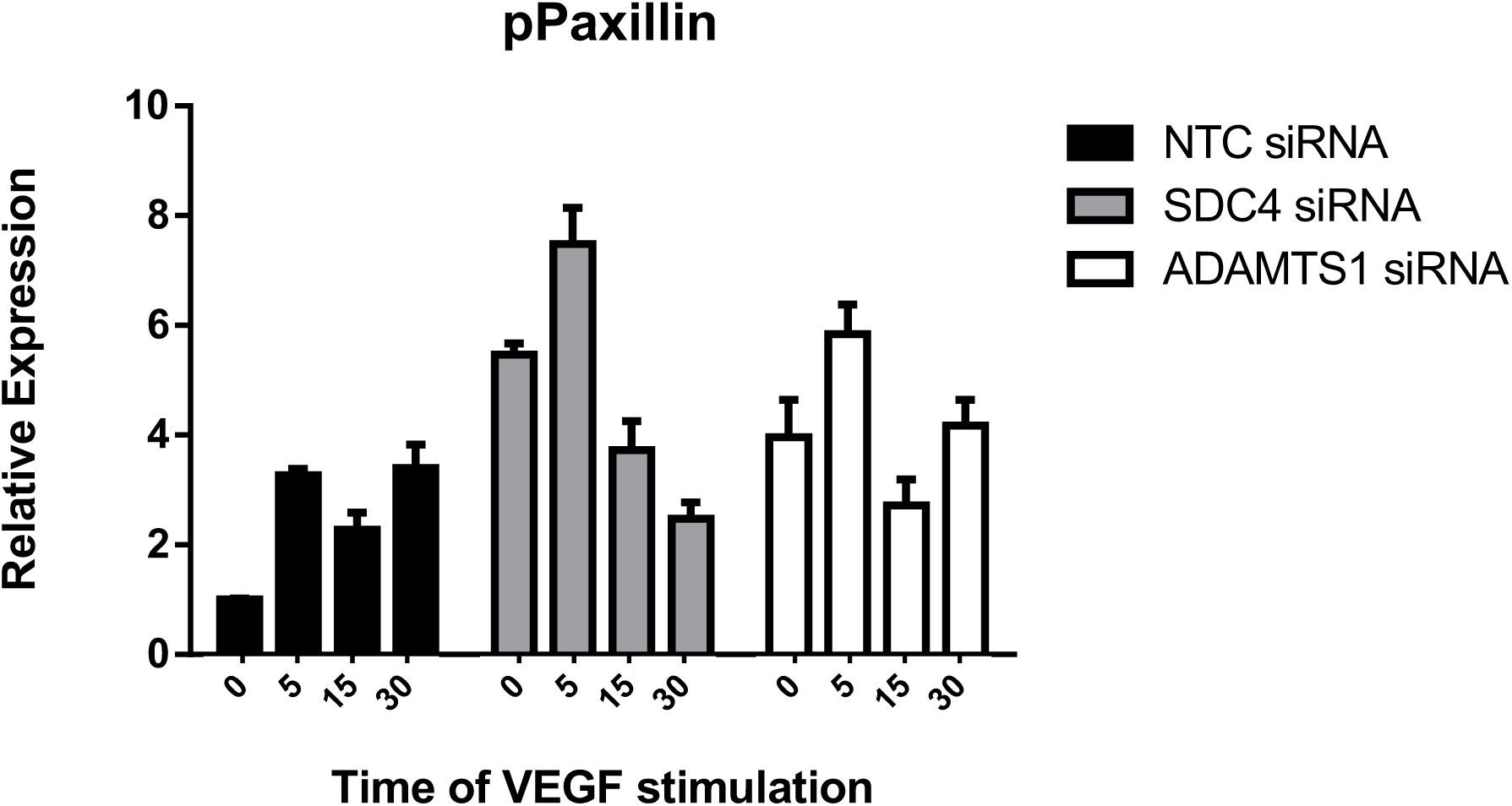
ADAMTS-1 or SDC4 depletion increases paxillin signalling in response to VEGFA. Image J densitometric quantification of VEGF timecourses (figure 5C). Phosphorylated paxillin expression level was calculated relative to total protein. (N=3).

**Figure S6:**
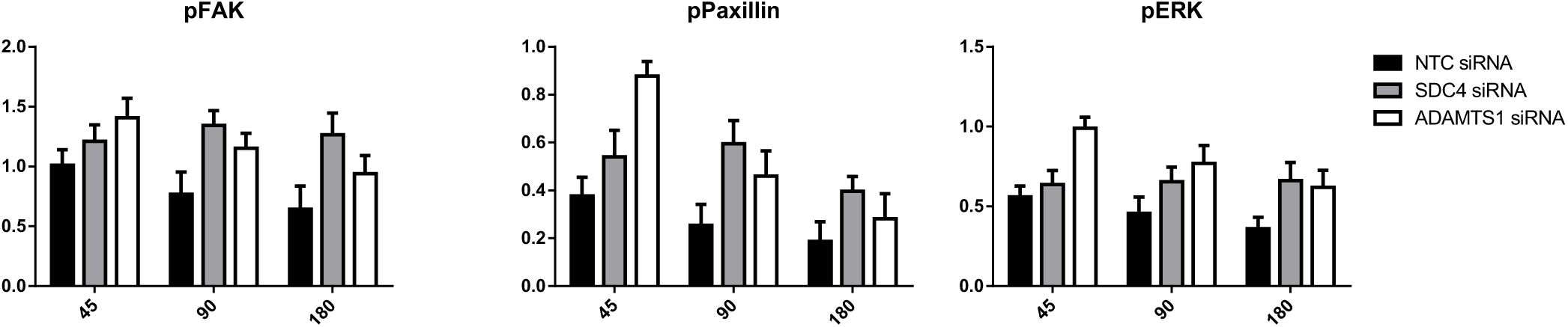
untreated cells show increased signalling responses to conditioned matrix. Image J densitometric quantification conditioned matrix timecourse (figure 5C). Phosphorylated protein expression levels were calculated relative to total protein. (N=3).

**Figure S7:**
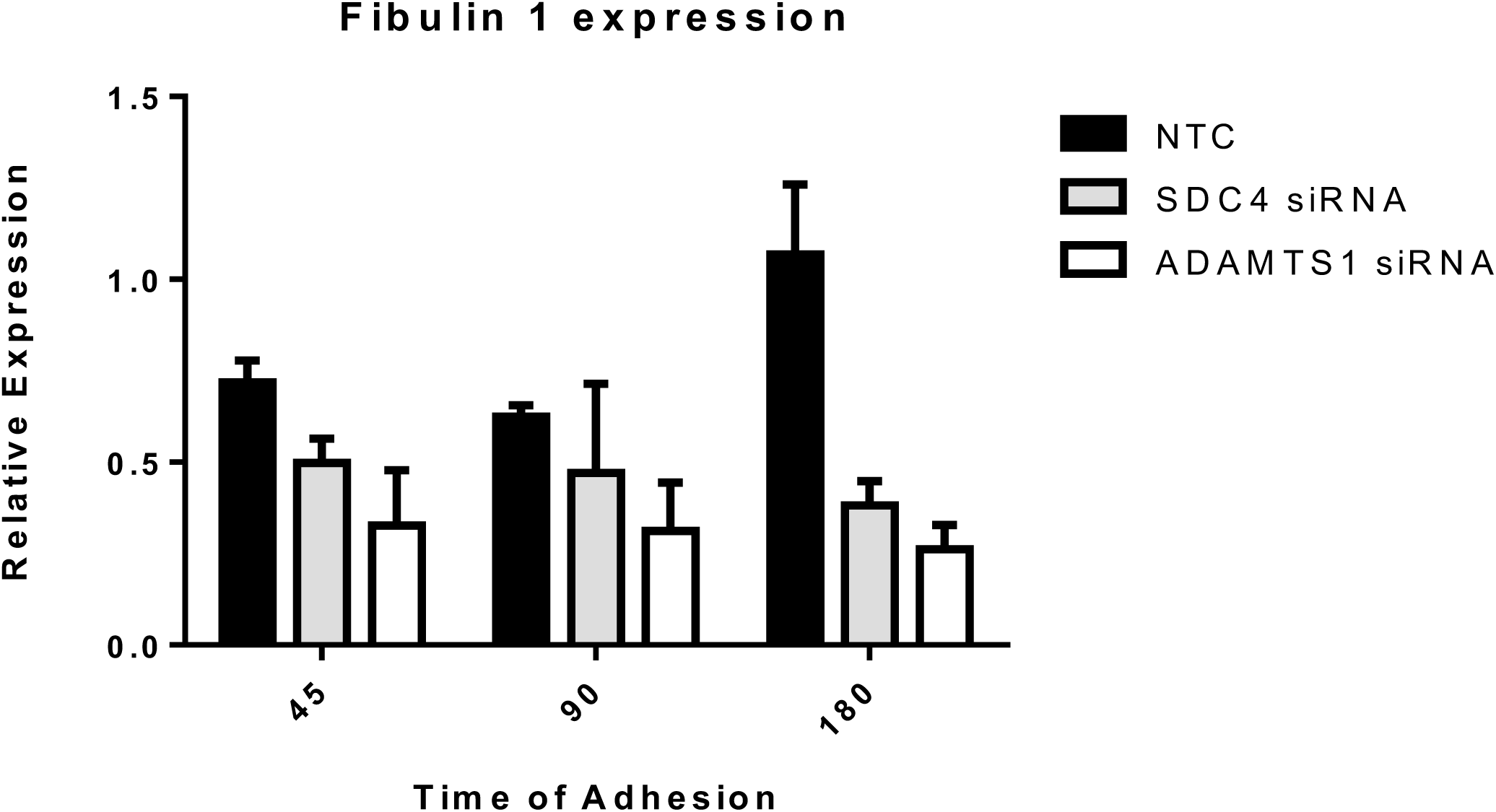
NTC cells, but not ADAMTS-1 or SDC4 siRNA treated cells, express increased fibulin 1. Image J densitometric quantification of fibronectin adherence timecourses (figure 7C). Fibulin 1 expression was calculated relative to GAPDH loading control. (N=3).

